# Inferring species compositions of complex fungal communities from long- and short-read sequence data

**DOI:** 10.1101/2021.05.02.442318

**Authors:** Yiheng Hu, Laszlo Irinyi, Minh Thuy Vi Hoang, Tavish Eenjes, Abigail Graetz, Eric Stone, Wieland Meyer, Benjamin Schwessinger, John P. Rathjen

## Abstract

**Background:** The kingdom fungi is crucial for life on earth and is highly diverse. Yet fungi are challenging to characterize. They can be difficult to culture and may be morphologically indistinct in culture. They can have complex genomes of over 1 Gb in size and are still underrepresented in whole genome sequence databases. Overall their description and analysis lags far behind other microbes such as bacteria. At the same time, classification of species via high throughput sequencing without prior purification is increasingly becoming the norm for pathogen detection, microbiome studies, and environmental monitoring. However, standardized procedures for characterizing unknown fungi from complex sequencing data have not yet been established.

**Results:** We compared different metagenomics sequencing and analysis strategies for the identification of fungal species. Using two fungal mock communities of 44 phylogenetically diverse species, we compared species classification and community composition analysis pipelines using shotgun metagenomics and amplicon sequencing data generated from both short and long read sequencing technologies. We show that regardless of the sequencing methodology used, the highest accuracy of species identification was achieved by sequence alignment against a fungi-specific database. During the assessment of classification algorithms, we found that applying cut-offs to the query coverage of each read or contig significantly improved the classification accuracy and community composition analysis without significant data loss.

**Conclusion:** Overall, our study expands the toolkit for identifying fungi by improving sequence-based fungal classification, and provides a practical guide for the design of metagenomics analyses.

## Introduction

Fungi are ubiquitous yet their presence and impact are often overlooked. It has been estimated that 2.2-3.8 million species inhabit planet earth [1] but only about 4% of these are catalogued [2]. Mora *et al*. estimated that there are 7.8 million and 298,0000 animal and plants species on earth with 12.3% and 72.4% of these characterised scientifically, respectively [3], which points towards a more central role in cultural awareness. In contrast, fungi are introduced to our consciousness via a brief mention in high school textbooks, or as largely side subjects in botany and microbiology courses at university [4,5]. Fungi play diverse roles throughout evolution and are particularly active in mediating the breakdown and uptake of nutrients. They constitute a major disease load to humans, causing millions of deaths per year, and wreak devastating crop losses via a constant toll of disease and epidemics and are an existential threat to many frog species [6,7]. On the other hand, fungi are or are used to manufacture delicious foods and beverages, and have saved countless lives via antibiotic production [8,9]. Therefore, a recent call was made to expand fungal research and improve our awareness of this special kingdom [10].

To progress our understanding of fungal biology we need to be able to classify more species more precisely. Fungi have been an independent kingdom since 1969 [11] with addition of further phyla in early 2000 [12–16]. Historically, its taxonomy was based on morphological and reproductive traits but this has been surpassed by DNA-based classification which revolutionized mycology, not only refining the conventional taxonomic tree [17,18] but also standardizing the identification of new species. In the absence of whole genome data, DNA-based classification primarily exploits the internal transcribed spacer (ITS) within the ribosomal RNA genes as a highly polymorphic marker to distinguish species. It is easily amplified and sequenced due to highly conserved flanking sequences and contains a high degree of variation between even closely related species. Although a mature pipeline comprising ITS amplification, IIllumina sequencing and data analysis has been established[19], several studies reported biases from the sequencing technology used and from unevenly amplified fungal marker regions [20–22]. Recently, novel strategies exploiting long-range amplification and long-read sequencing have been developed to improve these classifications [23,24]. In addition, whole genome shotgun sequencing and rapidly expanding genome databases allow mapping of newly generated DNA sequences directly to the database. This strategy allows exploitation of genetic variation throughout the genome and abandonment of the marker gene amplification step, which increases classification accuracy and reduces the biases from the estimation of relative abundance [25].

Although advanced sequencing methods allow novel strategies for fungal identification particularly from mixed samples, new demands are placed on data analysis pipelines to improve the accuracy of fungal classification. Different algorithms have been developed to classify DNA sequences at distinct taxonomic ranks based on sequence databases with taxonomic information [26–30]. For example, alignment algorithms such as Basic Local Alignment Search Tool (BLAST) [27] detect matches of each sequence to subjects of the target database along with the taxonomic information assigned to each entry. Alternatively, sequence features represented by short unique subsequences named k-mers can be derived from sequence data and mapped to databases to identify taxa with the highest number of cross-mapping k-mers[28]. Several studies have critically assessed algorithms for species classification on simulated datasets or bacterial community datasets [31–33], but comparisons of sequencing strategies for complex fungal communities alignment using real data and different identification pipelines are extremely rare. In addition to search algorithms, the choice of database also influences classifications dramatically, but only a few studies have researched their impact [34–36]. Therefore, more comprehensive benchmarking of both classification algorithms and databases are needed to optimise identification pipelines.

Here, we assessed different combinations of algorithms and databases during processing of both short- and long-read sequencing data for the identification of taxa from complex mock fungal communities. We identified key factors that influence the accuracy of classifications, both for mock community datasets and public datasets. Optimisation of these methods also lead to more accurate community composition analysis. Our results provide guidelines for the design of sequence-based community analysis for fungal species.

## Results

### Construction of mock fungal community datasets

We constructed two mock communities from the same set of 44 fungal species (Supplementary Table S1). Most of these are human-associated pathogenic yeasts while some are basidiomycete pathogens. One community comprised pooled DNA (PD) from each species and the second was composed of DNA extracted from equal quantities of fungal biomass (PB) of each species that were mixed together prior to extraction. We generated four sequence datasets for each community using Illumina and nanopore technologies, sequencing both shotgun metagenomes and targeted amplicons respectively. The data derived from each strategy are summarized in Table 1.

**Table 1.** The characteristics for each dataset.

| Sample | Sequencing Tech | Sequencing Strategy | # Basepairs | # reads | # Assembled contigs | # Mapped basepairs (Gb) |
| --- | --- | --- | --- | --- | --- | --- |
| PD | Illumina | Shotgun | 3.91 Gb | 14525058 | 338823 | 3.69 |
|  |  | Amplicon | 66.9/95.8/106.4 Mb<sup>a</sup> | 39374/9614/10236<sup>b</sup> | N/A | N/A |
|  | Nanopore | Shotgun | 1.96 Gb | 1273484 | N/A | N/A |
|  |  | Amplicon | 71.5/72.5/86.5 Mb | 26212/ 26680/ 31826<sup>b</sup> | N/A | N/A |
| PB | Illumina | Shotgun | 3.67 Gb | 13623120 | 345009 | 3.44 |
|  |  | Amplicon | 55.7/38.1/71.9 Mb<sup>a</sup> | 23613/13828/27093<sup>b</sup> | N/A | N/A |
|  | Nanopore | Shotgun | 3.78 Gb | 1043343 | N/A | N/A |
|  |  | Amplicon | 54.5/49.4/42.0 Mb | 20163/ 18273/ 15502<sup>b</sup> | N/A | N/A |

<sup>a</sup> The total basepairs of each technical replicate were calculated before importing into QIIME2 pipeline.
<sup>b</sup> Number of nanopore reads or paired-end Illumina reads for technical replicate 1/replicate 2/replicate 3 after quality control.

### Alignment algorithm against a specific fungal database resulted in the most accurate fungal classifications

We compared different analysis strategies for each shotgun dataset. For nanopore datasets, we directly used the quality-controlled reads for classification. For Illumina data, we quality filtered all reads and assembled them into contigs before classification to maximize the classification accuracy. We performed both alignment and k-mer based classifications on these data using BLAST and Kraken2 [27,29] using a ‘winner-takes-all’ strategy in which the top hit was taken as the identity of the query sequence. For each algorithm, we compared the use of two reference databases: the non-redundant NCBI nucleotide database (nt) [37] and the RefSeq fungi database (RFD) [38] which only contains curated fungal genomes. We first assessed the performance of each alignment tool on both databases for each data input. We compared the concordance in the results of each pipeline at the genus level. We define concordance as the percentage of fungal genera identified by both analyses in a pairwise comparison (Figure 1A). The concordance between analyses on each dataset varied between 69% and 86% and generally, Illumina data resulted in higher concordance than did nanopore data.

**Figure 1.**
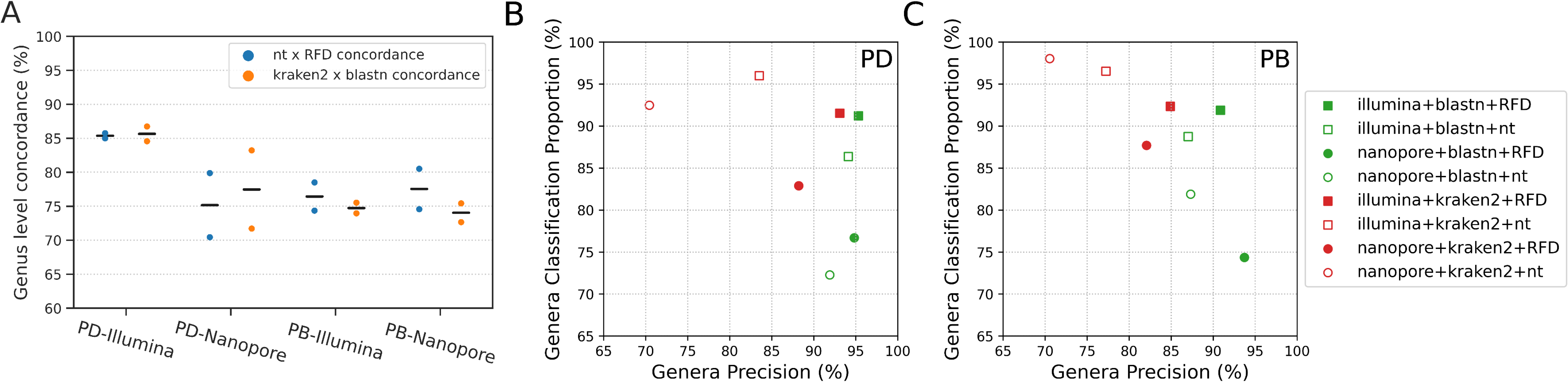
Analysis of shotgun metagenomics data. (A) Swarmplot showing the concordance in genus identification after varying either the alignment algorithm or querying different databases on different data inputs. nt = NCBI nucleotide database; RFD = RefSeq Fungi database; data inputs are indicated below the line (PD = pooled data; PB = pooled biomass); (B) Identification of fungal genera from PD samples. The classified proportion and precision were derived from different combinations of search algorithms and databases as indicated (box); (C) Identification of fungal genera from PB samples. The classification proportion and precision were derived from the different combinations of search algorithms and databases as indicated.

We then aimed to identify the combination of algorithm and database that yielded the most accurate species identification. We used classified proportion and precision to evaluate each classification, where 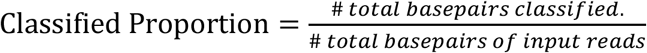, and 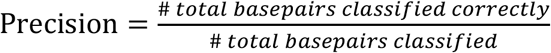. The number of total basepairs is calculated as total read length for nanopore reads and total coverage of Illumina reads to each contig [32,33]. We plotted the precision and classified proportion for each pipeline and found three regular patterns (Figures 1B and 1C): First, for each dataset, BLAST resulted in higher precision but lower classification proportion by comparison to Kraken2. Second, Illumina contigs returned higher classification proportion and precision than nanopore reads. Third, classification against the RFD database yielded higher precision than those against the nt database. In summary, BLAST alignments against the RFD database yielded the best classification strategy.

### Applying cut-offs to query coverage improves classification accuracy on shotgun metagenomics datasets

We next aimed to improve our classification scheme by filtering the BLAST search results. We reasoned that restricting alignment metrics would reduce the number of false classifications. To investigate changes in classification accuracy after restricting BLAST output parameters, we first BLASTed shotgun metagenomics reads against the RFD database without applying any filter, then applied progressive cut-offs on different parameters of the BLAST results. We evaluated changes in the results based on the metrics precision, remaining rate and completeness. Precision is described above and estimates the accuracy of the classification; remaining rate captures the percentage of the input data remaining after the application of each cut-off; and completeness is the number of taxa captured relative to the total number of taxa within the mock community. We initially applied cut-offs on query length and two alignment metrics; E-value - the number of expected hits of similar quality that could be found by chance alone; and pident – the percentage of identical matches within the region of alignment between query and subject. As shown in Figure 2A, applying progressive cut-offs to query length did not improve the precision, whilst both completeness and remaining rate diminished dramatically from very small cut-off values. Cut-offs applied to alignment E-values removed <20% of the BLAST results, whereas precision showed minor improvement, especially on nanopore datasets (Figure 2B). For Illumina data, applying cut-offs to the E-value increased the precision by around 2% but at the cost of diminished completeness. E-value cut-offs performed better on nanopore datasets, improving precision by 3% (PD) or 4% (PB) with non-identification of only a single genus from the mock community, at 10^−250^ or almost 10^−400^ respectively. Progressive cut-offs on pident yielded the best results of all three filters. For Illumina data, precision was improved by up to 8% for PB data, and completeness remained at 100% in almost all cases (Figure 2C). For nanopore datasets, pident cut-offs improved the precision by up to ~3% before sharp decreases, with a concurrent filtering of ~60% BLAST result as shown by the remaining rate. Given the characteristically high error rate of nanopore reads, we also applied cut-offs on quality scores to these data. Cut-offs applied to Phred scores did not alter the precision, while a significant proportion of the dataset was lost through filtering (Supplementary Figure S1). Overall, our results suggest that applying each filter to BLAST results performs well on either Illumina or nanopore data but not both, and that cut-offs based on query length or quality scores did not affect the precision significantly.

**Figure 2.**
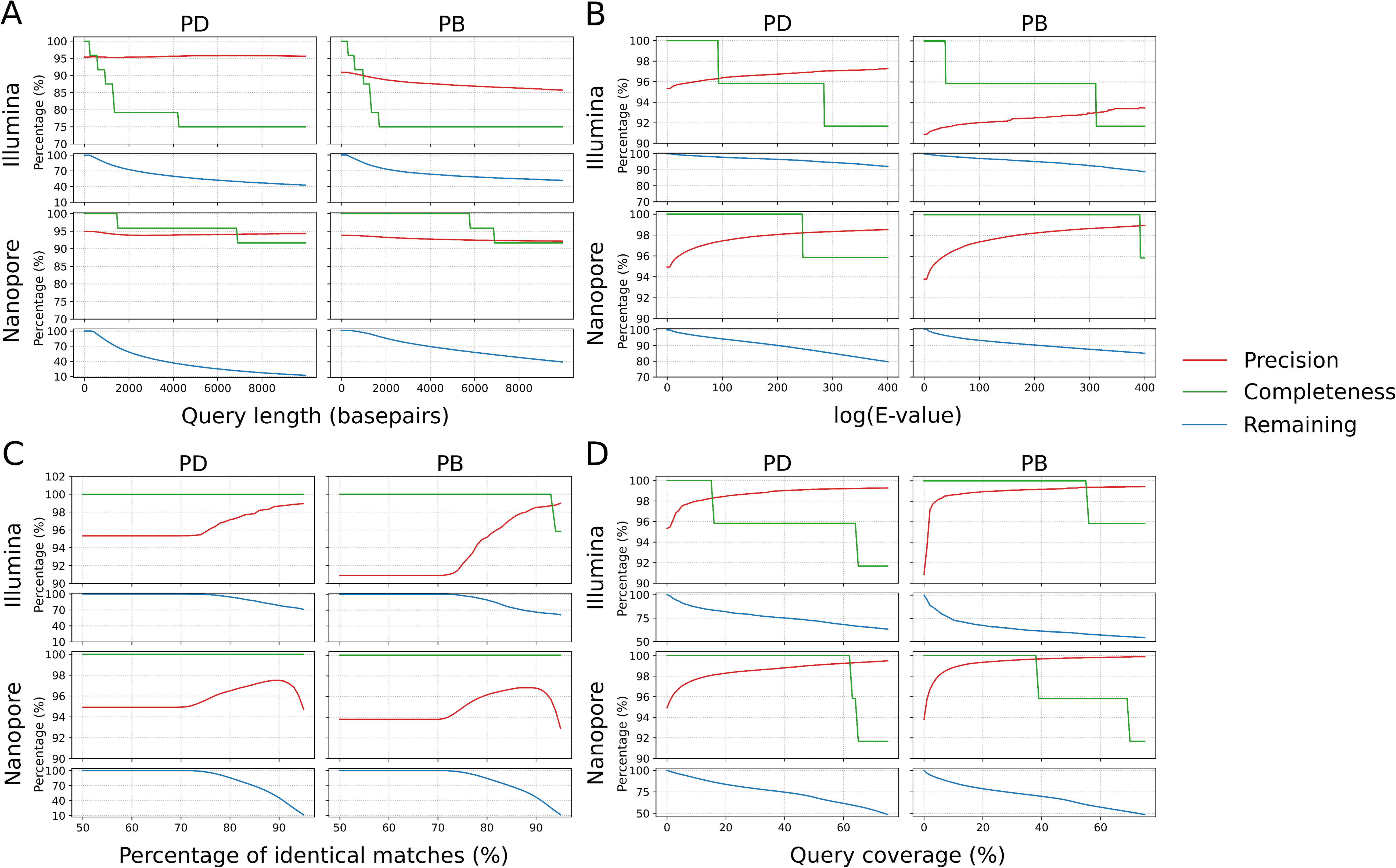
Dynamics in precision, completeness and remaining rate after applying progressive cut-offs on BLAST alignment metrics. (A) Cut-offs applied to query length. (B) Cut-offs applied to alignment E-values. (C) Cut-offs applied to the percentage of identical matches. (D) Cut-offs applied to query coverage.

Given the results above, we investigated how the alignment parameters were calculated and explored other variables to improve the classifications. The BLAST E-value is calculated as E = mn2^−S^ in which S is the bits score derived from the number of gaps and mismatches in the alignment, and m and n are the query length and database total length respectively [39]. Therefore, the E-value is influenced exponentially by the alignment quality. We next investigated query coverage, a metric based on how much of the query sequence aligned to the subject. We calculated the query coverage as the number of identical matches divided by the read or contig length, and applied progressive cut-offs on this parameter for each dataset/algorithm analysis. As shown in Figure 2D, applying cut-offs on query coverage improved the precision of all four analyses significantly, and did not cause losses of completeness at smaller cut-off values. For example, at a 10% cut-off on query coverage, the precision of all four analyses was 98-99% while the completeness remained at 100% and the removed BLAST results ranged from 10-25%. This result not only supported our hypothesis that the total length of the alignment matters as much as the alignment quality, but also suggested a novel approach to improve the accuracy of fungal classification.

### Improving taxa identification from published metagenomics datasets using query coverage as a filtering parameter

After improving classifications by applying cut-offs to the query coverage on the mock community datasets, we extended this strategy to try to improve the classification of published shotgun metagenomics datasets. We re-analysed ten nanopore and six Illumina shotgun metagenomics datasets [40–43]. These included host-associated fungal samples (nanopore) and host-depleted microbiome data (Illumina). Since the environmental datasets contain unknown species, we followed the concept of classification precision. We calculated the percentage of the dataset that was classified into taxa known to be included in the sample. For example, in re-analysing human clinical samples [42], we included the pathogen (*Pneumocystis*) and the host human (*Homo*) as the true taxa, and calculated the total proportion of query sequences classified to these taxa before and after applying cut-offs on query coverage. Table 2 shows the improvement in taxonomic classification from the published datasets after applying query coverage cut-offs. We initially applied a 20% cut-off on the query coverage for all analyses, but the data loss in most cases was too high. Therefore, we applied query cut-offs that filtered around 20% of the blast result based on our results from the mock fungal community datasets (Figure 2D).

**Table 2.** Assignment of published sequence data to genera after application of cut-offs to query coverage.

| Sample ID | Sample description | Sequencing tech | Cut-offs on query coverage (%) | Filtered results (%) | Percentage of confirmed genera **BEFORE** applying cut-offs (%) | Percentage of confirmed genera **AFTER** applying cut-offs (%) |
| --- | --- | --- | --- | --- | --- | --- |
| a1 | Human sputum samples<sup>42</sup> | Nanopore | 59 | 20.2 | 85.9 | 86.5 |
| a2 |  |  | 53.2 | 20.1 | 97.9 | 98.5 |
| a3 |  |  | 54 | 20.5 | 96.5 | 97.4 |
| a4 |  |  | 45.5 | 20.1 | 16.2 | 19.8 |
| a5 |  |  | 58.5 | 20 | 71.1 | 66.9 |
| a6 |  |  | 50.4 | 20.1 | 93.6 | 94.7 |
| b1 | Field infected wheat samples<sup>41</sup> |  | 5 | 20 | 60.4 | 75.1 |
| b2 |  |  | 0.77 | 19.9 | 34.8 | 43 |
| b3 |  |  | 12 | 19.7 | 67 | 82 |
| b4 |  |  | 0.61 | 20 | 5.8 | 6.2 |
| c1 | Pig gut microbiome samples<sup>43</sup> | Illumina | 2.4 | 20.1 | 32 | 35.4 |
| c2 |  |  | 3.3 | 20.2 | 34.2 | 36.6 |
| c3 |  |  | 2.6 | 20.2 | 35.2 | 38.3 |
| d1 | Mouse gut microbiome samples<sup>40</sup> |  | 3.4 | 19.8 | 29.1 | 24.3 |
| d2 |  |  | 14 | 20.1 | 63.7 | 69.4 |
| d3 |  |  | 4.5 | 20.2 | 38.6 | 42.3 |

For all Illumina datasets, we downloaded the quality-controlled sequences and re-analysed them using the assembly and BLAST pipeline described above against the NCBI nt database. For the nanopore human datasets [42], we used the BLAST results taken directly from the original articles for analysis. For the infected wheat datasets [41], we downloaded the sequences and re-analysed them against the RefSeq fungal database. The precision increased for nearly all datasets after applying query coverage cut-offs (Table 2). For the Illumina microbiome datasets, we first assessed the change of proportions in fungal taxa after applying cut-offs on query coverages using the species lists identified by Donovan *et al.* [44] as confirmed taxa. We observed only marginal increase in percentages for the confirmed fungal communities, due to their low total proportions in the original samples. We then calculated the improvement in precision for the bacterial communities. The Illumina datasets were generated from swine and mouse gut microbiome samples, so we assessed the change in proportions of their core bacterial genera (a group of bacteria commonly present in swine and mouse guts [45,46]). The percentages of confirmed core bacterial genera improved by up to 5.7% after applying cut-offs on query coverage (Table 2). In addition, in the nanopore human datasets, the total percentage of reads classified as *Homo* in the three healthy individual samples were improved by applying cut-offs to query coverage. These results indicated that this strategy may be broadly applicable not only to fungal species, but also to the classification of other eukaryotes and bacteria. One Illumina dataset (d1) and one nanopore dataset (a5) showed decreased percentages of confirmed taxa after applying query coverage cut-offs, which might be because the core microbiome species are not representing the species identified in the Illumina sample, or due to the low coverage and high error rate of nanopore data.

### Benchmarking classification pipelines for amplicon datasets identified advantages of each strategy

We next assessed different strategies for the classification of ITS amplicon datasets. We amplified the ITS region from both mock communities using two different primer pairs and three technical replicates for each sample. Taking advantage of nanopore technology, we performed long-amplicon sequencing of a roughly 3 kb ribosomal RNA gene region covering part of the 28S subunit, ITS1, 5.8S subunit, ITS2 and part of the 18S subunit [19]. For Illumina sequencing we used the well-established ITS1F-ITS2 amplicon of about 300 bp in length [47]. Similar to the analysis of the shotgun datasets, we applied both k-mer and alignment-based approaches to the classification of nanopore amplicon data. We used the pair-wise alignment algorithm minimap2 as the alignment algorithm instead of BLAST due to its speed and efficiency. We tested four different databases for classification of long amplicons; the NCBI 18S and 28S databases, and two ITS databases from NCBI and UNITE, respectively [38,48]. Overall, we found that the k-mer algorithm returned much higher classification proportion than alignment for each nanopore dataset, but the highest precision (~97%) were achieved by combining the minimap2 alignment algorithm with the NCBI ITS database (Figure 3A). For Illumina amplicon datasets, we applied the QIIME2 pipeline which is one of the most widely used strategies for ITS classification and community composition analysis[49]. The QIIME2 pipeline groups similar Illumina amplicons into sequence features before classification to reduce the demand on computational resources [50]. Since all individual Illumina reads are grouped into sequence features and all the sequence features are classified, the classification proportion of the Illumina amplicon datasets are 100%. We plotted precision rates from the QIIME2 analysis of both the PD and PB samples with their means (Figure 3B). The mean precision from either Illumina dataset were lower than that from k-mer analysis of the respective nanopore datasets.

**Figure 3.**
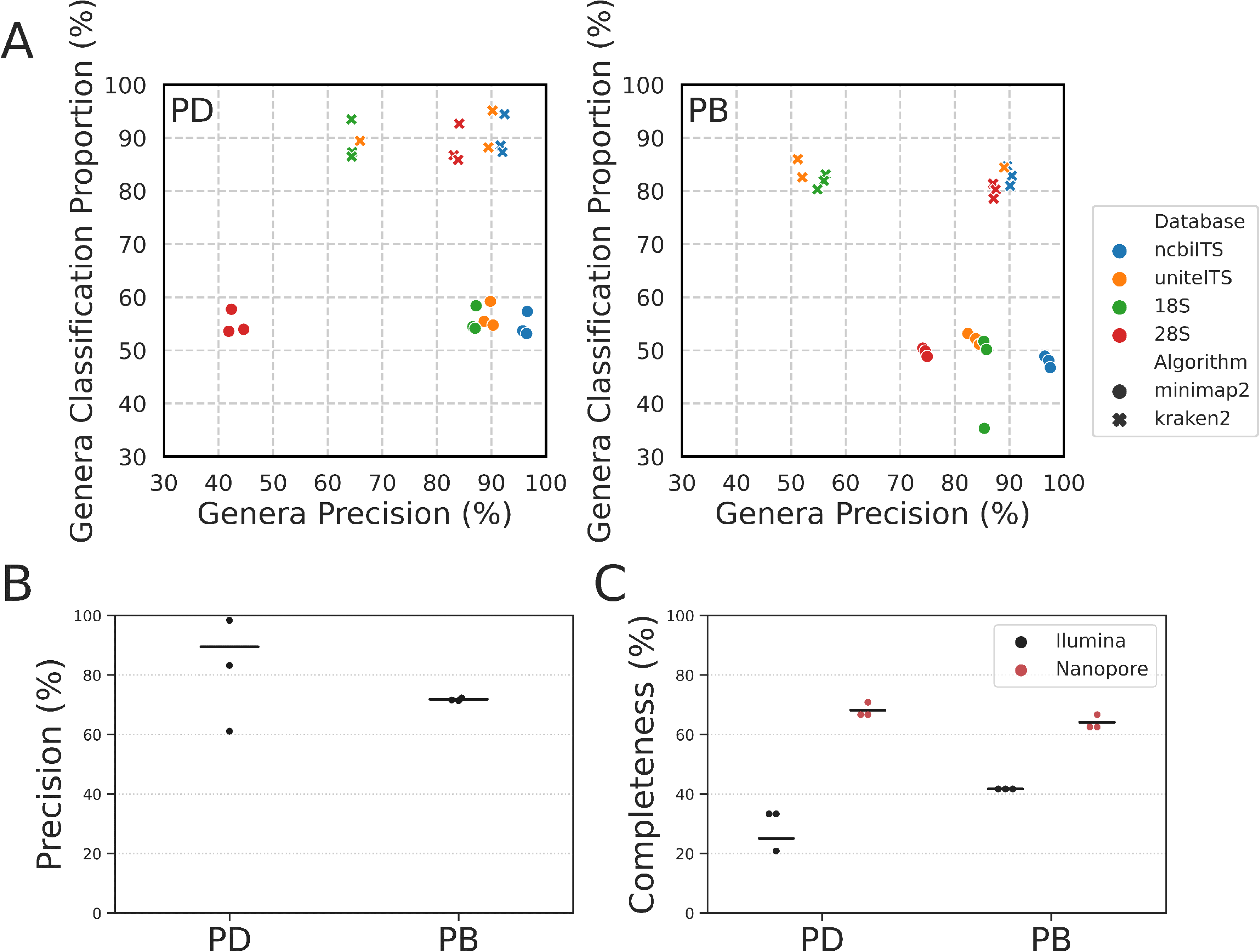
Benchmarking of amplicon datasets. (A) Scatter plot represented genus level classification proportion and precision for nanopore amplicon data. (B) Genus level precision of Illumina amplicon data. Classification proportion of Illumina data were 100% due to the nature of the QIIME2 pipeline (based on the UNITE ITS database). (C) Genus level completeness of both nanopore and Illumina amplicon datasets. The nanopore results are from minimap2 algorithm and uniteITS database.

Although the precision from the amplicon datasets were higher than that from shotgun datasets, the ITS classification did not identify all genera within the mock community, as shown by our completeness analysis (Figure 3C). The nanopore amplicons identified 68% (PD) and 63% (PB) of the total genera in the mock community, whereas the Illumina amplicon datasets covered only 25% and 41% of the genera respectively. We suspect that the low completeness from ITS classifications was due partially to the low quality of this particular dataset (Table 1) and partially due to non-uniform amplification from the different primer pairs. However, there were fewer nanopore amplicon reads than in the Illumina amplicon datasets and the completeness from the nanopore data was higher (Figure 3C). This supports the argument that long amplicons identify a wider range of species and are more accurate in species classification than short amplicons [51,52].

### Cut-offs on query coverage also improve community composition analysis

We next analysed community compositions using the most accurate classification method for each dataset. Community composition refers to the identity and relative abundances of all taxa in a community. Given the observation that use of a restricted database resulted in higher classification precision, we constructed a database containing only the genomes from species within the mock community and aligned all of the data to the mock community database using BLAST. This forces the precision to 100% as any classification will belong to a species from the mock community. We then BLASTed each dataset against this database and calculated the relative abundance of each genus. We defined this as the ‘gold standard’ for community composition analysis of the mock fungal community (Figure 4A). We then compared the community composition determined from each combination of algorithms and databases with the gold standard for each dataset, and measured their differences using three statistical distance tests: Bhattacharyya distance, relative Euclidean distance and relative entropy [53–55]. Consistently, BLASTing sequences against the RFD database produced community compositions with the highest similarity to the gold standard analysis (Figure 4B).

**Figure 4.**
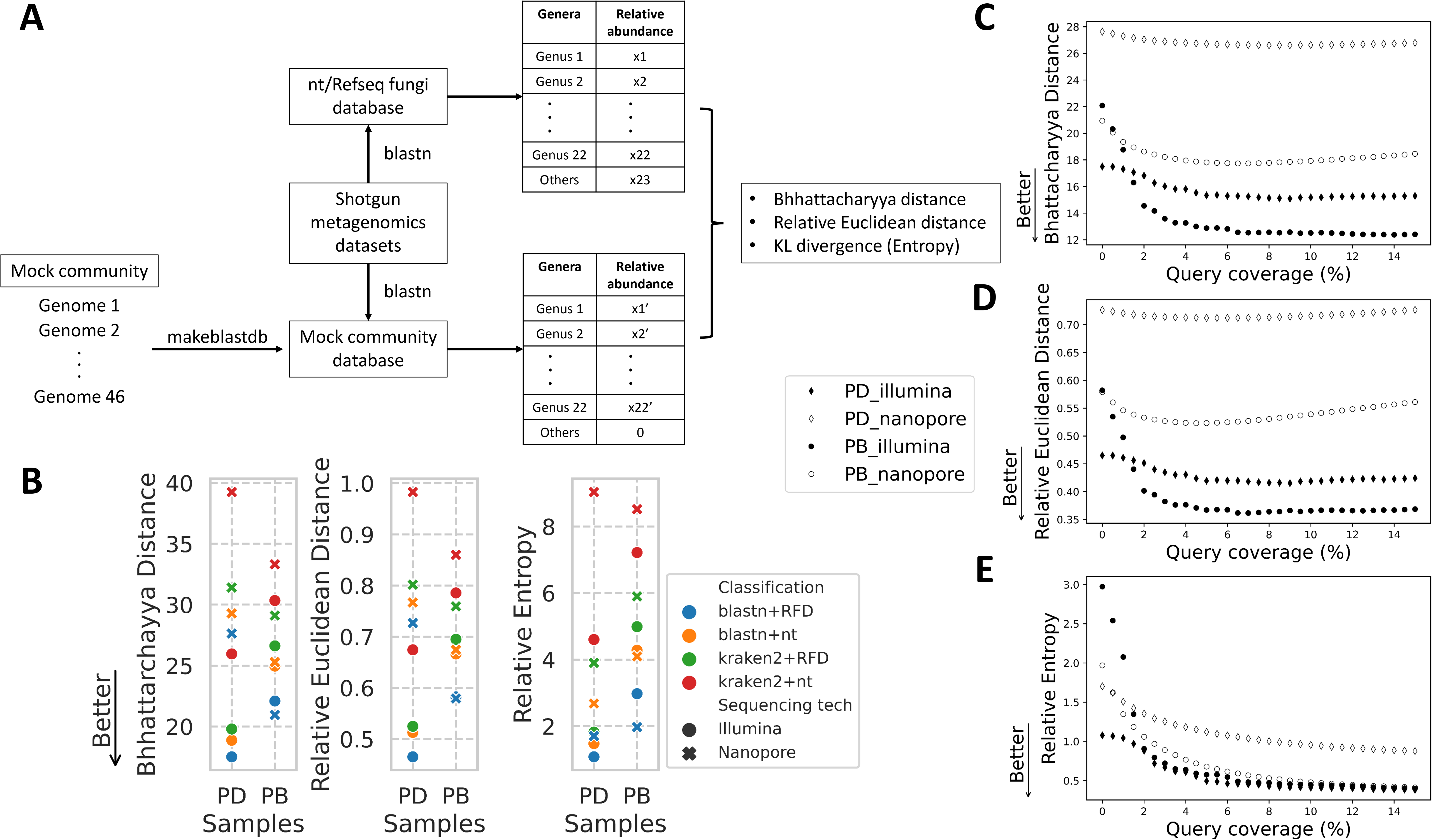
Improving community composition analysis by applying query coverage cut-offs. (A) Experimental flowchart for analysing community compositions. (B) Statistical similarity measures between gold standard community composition and each combination of algorithms and databases. Lower values correspond to greater similarity between the samples and the gold standard. (C) Change in Bhattacharyya distance after applying cut-offs to query coverage for each dataset as indicated. The query coverage gap between each dot point is 0.5%. (D) Change in relative Euclidean distance after applying cut-offs to query coverage for each dataset. The gap between each dot point is 0.5%. (E) Change in relative entropy after applying cut-offs on query coverage for each dataset. The gap between each dot point is 0.5%.

To assess whether query coverage cut-offs also improved the community composition analysis of shotgun metagenomics data, we plotted the changes in statistical distance after progressive application of query coverage cut-offs (Figure 4). After applying cut-offs on the query coverage, the community composition improved in all cases especially for lower cut-off values. The community compositions from PB-Illumina datasets improved and turned out to be the most similar to the gold standard at query-coverage cut-offs greater than 3 - 4%, which is consistent with the changes in precision rate shown in Figure 2D. Overall, our results illustrated that applying cut-offs on query coverage did not only improve the classification accuracy, but also the community composition analysis.

## Discussion

Here we investigated the taxonomic classification from sequencing data, one of the key steps in all metagenomic workflows, with a particular focus on fungi. After assessing various combinations of algorithms and databases following different sequencing strategies, we found that the combination of BLAST with the specific RFD database always resulted in the most precise classification for all mock fungal community datasets. These classifications were further improved when applying cut-offs on query coverage including positive flow on effects on downstream community composition analysis from shotgun metagenomics datasets.

Despite that sampling and DNA extraction substantially influence the outcome of species classifications [56–58], choosing an appropriate sequencing strategy is the primary step towards accurately profiling a sample. For shotgun datasets, our results suggested that both short and long shotgun datasets have comparable accuracy and both higher than the amplicon datasets. However, Illumina shotgun datasets require additional steps to assemble reads into contigs before querying them against a database, and to map all reads back to the assembly to quantify the coverage. These processes are necessary to achieve accurate classification from longer contigs [59], but result in a longer sequence-to-result turnover than the long read shotgun data. In the analysis of the amplicon data, long range amplicons performs better in the classification accuracy and completeness compare to the short ITS data, consistent with other studies [51,52]. Comparing to the results from shotgun datasets, the overall completeness from the result of amplicon datasets is much lower. We think that is because we used much less amplicon data for benchmarking classification pipelines, and the incomplete database which do not contain all taxa present in our mock community. Overall, the long read shotgun datasets returned the most accurate fungal classification.

Next, our data supported that alignment algorithm (BLAST) outperform the k-mer based approach (kraken2) in the accuracy of classification [32,60], and also compared progressive cut-offs to major alignment parameters for shotgun metagenomics data. We found that applying read length or read quality cut-offs did not improve the precision of the classification for all shotgun datasets. This observation is different with the previous study based on simulated data, which claimed that the long reads improves the accuracy of classification [60]. Cut-offs on pident slightly improved the classification accuracy for illumina datasets, but the error-prone nature of the nanopore data (~10% error rate) is also reflected in the result, as it causes the breakdown of precision when pident cut-offs reach 90% (Figure 2C).

We found that query coverage cut-off that filter out 20% blast result worked best. Unlike the E-value weighing the gaps and mismatch as the major factor effecting alignment quality, the query coverage weighs the query length as well as the number of identical matches in the assessment of the alignment quality. In this case, we can eliminate more spurious alignments that are due to a small proportion of reads with high fidelity to the reference, which are commonly found in reads containing conserved genes and repeated sequences. Interestingly, to reach the same 20% filtering threshold, we set up higher cut-offs on the query coverage (10-20%) in mock community datasets than the real environmental datasets, including few extremely low thresholds of query coverage in the Illumina shotgun datasets. We compared other studies that use simulated data to generate metagenomics contigs for classification, and found that they used 90% query coverage cut-offs as the parameter[62–64]. Together with the different result of read length and read quality cut-offs, this observations highlighted the difference between the use of real environmental data and the simulated data in benchmarking studies, especially for the classification of complex microbial communities.

PD and PB samples showed slightly different results in comparing statistical distance with the gold standard. After applying cut-offs on query coverage, both Bhattacharyya distance and Euclidean distance between the best practice and the gold standard classification only showed marginal decrease in PD samples, and slowly reversed as the cut-offs increase. We think that is because about 1/3 of reads were classified as *Candida* in the pooled DNA sample, so the difference on the relative abundance of one *Candida* genus between the gold standard community composition and the best practice is much higher and much more influential to the final distance than that from other genera.

Following the importance of the alignment quantity represented by the query coverage, the next question is, how to bring the low quality but high quantity alignment into consideration? Therefore, the winner-takes-all selection strategy itself can be re-designed, as the highly conserved genome regions from different species generate highly close alignment scores between the best alignment and other top alignments. In this case, a weighing statistics and the relative probability for multiple top taxonomic assignments can be explored and introduced to replace the best-hit-takes-all strategy. This will be particularly useful in connection with the rapid expansion of the fungal genome databases.

Next to the right classification tool, chosen the appropriate database significantly influences analysis outcomes [33,34]. Based on our observation, we suggest that ‘prior knowledge’ about the dataset should guide the choice of the appropriate database as this will improve the accuracy of taxonomic classifications. For example, our results suggested that the restricted database resulted in more accurate fungal classifications for shotgun metagenomics datasets. This strategy might be appropriate if queries are initial binned into kingdoms before a more in-depth analysis with kingdom specific databases. Also, Kaehler *et al.* [65] incorporated environment-specific taxonomic abundance information into the analysis of amplicon datasets and showed that these improve classification accuracy. Similar approaches can be applied to metagenomic datasets. In addition, machine learning strategies become increasingly popular for analysing genomic data. Here taxonomic classifiers could be trained on existing labelled sequence datasets before being applied to communities with similar composition to the training datasets or to identify target species from complex communities [60,66].

## Conclusion

In this study, we perform an in-depth analysis on how different sequencing strategies, classification algorithms and databases impact fungal classifications using complex real-life mock community sequencing datasets. We find that alignment algorithm (BLAST) with targeted fungal database (RFD) achieve the best classification accuracy and community composition estimates. These can be further improved by applying cut-offs on query coverage. Taken together, the findings from our benchmarking workflows have important implications for mycology studies for multiple stages of metagenomics analysis, and provided a guide to other researchers aiming to study fungal metagenomics.

## Methods

### Code availability

All detailed commands and scripts used in each step were summarized in https://github.com/Yiheng323/Benchmarking-taxonomic-classification-strategies-using-mock-fungal-communities.

### Fungal harvesting, DNA extraction and construction of mock communities

Selected fungal strains were cultured onto Sabouraud dextrose agar and incubated for 48 hours at 27°C.

For the species in the PD community, an inoculating loop full of fungal cells were scraped into a 1.5 mL microfuge tube and crushed with a pestle and liquid nitrogen. Genomic DNA was then extracted using the Zymo Research *Quick*-DNA Fungal/Bacterial Miniprep Kit (cat. no. D6005 Zymo Research, Irvine, CA, USA). First, BashingBead™ Buffer was added to the crushed fungal cells and vortexed. The mixture was then filtered through a Zymo-SpinTM III-F Column and the filtrate was combined with Genomic Lysis Buffer. The mixture was filtered through a Zymo-Spin™ IICR Column and washed with DNA Pre-Wash buffer and g-DNA Wash Buffer. The DNA was eluted in nuclease free water. DNA concentration was measured using the DeNovix dsDNA Broad Range Kit (DeNovix, Wilmington, DE, USA) and 250 ng of DNA from each strain were then pooled together.

For the PB community, two inoculating loops of fungi of each species in teg mock community were scraped into a ceramic mortar. Liquid nitrogen was then poured into the mortar and the fungal mixture was crushed into a fine powder. DNA was then extracted using the Qiagen DNeasy PowerMax Soil Kit (cat. no. 12988-10 Qiagen, Hilden, Germany). PowerBead Solution and Solution C1 were added to the crushed fungal community, vortexed and centrifuged. The supernatant was then added to Solution C2, mixed and centrifuged, which was then repeated with Solution C3. The resulting supernatant was combined with Solution C4 and centrifuged through a column. The column was then washed twice with Solution C5. Final DNA was eluted in nuclease free water and the concentration measured using the DeNovix dsDNA Broad Range Kit.

### Library preparation and sequencing

The ITS1 regions of the rRNA gene were amplified with the universal fungal primers, ITS1F (CTTGGTCATTTAGAGGAAGTAA) and ITS2 (GCTGCGTTCTTCATCGATGC)[47]. Sequencing of PCR amplicons was conducted with MiSeq^®^ System of Illumina (Illumina, San Diego, CA, USA) by the Australian Genome Research Facility. The Illumina bcl2fastq 2.18.0.12 pipeline was used to generate the sequence data. Pair-ends reads 2 × 300bp were generated up to 0.15 GB per sample for amplicon data. The Illumina amplicon data are then directly imported into QIIME2 for analysis. For shotgun Illumina datasets, we employed the same sequencing pipeline as the amplicon data, with MiSeq^®^ and bcl2fastq 2.18.0.12 pipeline for the 2 × 300bp paired end reads. Raw shotgun Illumina reads were trimmed adapters with Trimomatic [67]. Quality controlled, paired end reads were merged and assembled to metagenomics contigs using IDBA_UD [68], which is more suitable for datasets with uneven sequencing depths of each species. After assembly, raw reads were mapped back to the contigs using bwa-mem [69], and the bam files were generated and sorted from sam files using samtools [70]. Bedtools [71] was used for generating coverage for each contig, and we used python numpy and pandas module to calculate the average coverage for each contig.

For Nanopore sequencing of both shotgun and amplicon sequencing, we used Ligation Sequencing 1D SQK-LSK108 and Native Barcoding Expansion (PCR-free) EXP-NBD103 Kits from ONT (UK), as adapted by Hu and Schwessinger [72], which was adapted from the manufacturer's instructions with the omission of DNA fragmentation and DNA repair. DNA was first cleaned up using a 1× volume of Agencourt AMPure XP beads (cat. no. A63881, Beckman Coulter, Indianapolis, IN, USA) following manufacturer’s instructions. We then eluted the beads binded DNA in 51 μl nuclease free water and quantified using NanoDrop^®^ and Quibit™ Fluorometer (Thermo Fisher Scientific, Waltham, MA, USA). DNA was end-repaired (NEBNext Ultra II End-Repair/dA-tailing Module, cat. No. E7546), 1x volume beads cleaned (AMPure XP beads) and eluted in 31 μl nuclease free water. Barcoding reaction was performed by adding 2 μl of each native barcode and 20 μl NEB Blunt/TA Master Mix (cat. No. M0367) into 18 μl DNA, mixing gently and incubating at room temperature for 10 minutes. A 1× volume (40 μl) Agencourt AMPure XP clean-up was then performed and the DNA was eluted in 15 μl nuclease free water. Ligation was then performed by adding 20 μl Barcode Adapter Mix (EXP-NBD103 Native Barcoding Expansion Kit, ONT, UK), 20 μl NEBNext Quick Ligation Reaction Buffer, and Quick T4 DNA Ligase (cat. No. E6056) to the 50 μl pooled equimolar barcoded DNA, mixing gently and incubating at room temperature for 10 minutes. The adapter-ligated DNA was cleaned-up by adding a 0.4× volume (40 μl) of Agencourt AMPure XP beads, incubating for 5 minutes at room temperature and resuspending the pellet twice in 140 μl ABB provided in the SQK-LSK108 kit. The purified-ligated DNA was resuspended by adding 15 μl ELB provided in the SQK-LSK108 kit and resuspending the beads. The beads were pelleted again and the supernatant sequencing library was transferred to a new 0.5 ml DNA LoBind tube (Eppendorf, Germany). Nanopore sequencing was carried out by MinION MK1b device using R9.4.1 Flowcells. Raw fast5 files are barcode demultiplexed by deepbiner (ONT), then basecalled by Guppy (v3.6.0, ONT, UK). Quality passed reads in fastq files were trimmed adapters and barcodes using qcat (ONT, UK). For the long amplicon data, we filtered out reads less than 2000 base pairs. All sequencing data was submitted to NCBI Short Read Archive (SRA) under the bioproject PRJNA725368 including eight accessions: SRX10705648, SRX10705649, SRX10705650, SRX10705651, SRX10705695, SRX10705696 and SRX10705697.

### Genome assembly

While generating the reference genome database, we found that there were no reference genomes for *Candida rugosa*, *Candida mesorugosa* and *Cryptococcus magnus*, so we performed nanopore sequencing on pure DNA from each species and assembled their draft genomes. These assemblies were of sufficient contiguity and quality (Supplementary Table S2), so we added the new draft genomes into the reference database.

The nanopore data of *Candida rugosa*, *Candida mesorugosa* and *Cryptococcus magnus* was generated individually using Ligation Sequencing 1D SQK-LSK108 kit alone, and from independent flowcells. Data from each flowcell was basecalled and quality filtered using the same pipeline as described above. We got roughly 40X coverage for *Candida rugosa* and *Candida mesorugosa*, and 20X coverage for *Cryptococcus magnus.* Draft genomes were assembled with Flye [73] using default parameters and an estimated genome size of 20Mb. After assembly, the contigs were polished ten times with Racon [74] using nanopore reads, followed by one polishing with Medaka (ONT). Polished assembly was assessed completeness using BUSCO [75]. The assembly statistics were reported from Flye.

### Database constructions

For shotgun metagenomics analysis, we used three BLAST database and three kraken databases. Two databases (nt and RFD) are from the same NCBI source, downloaded in May 2019. BLAST and kraken2 nt databases were downloaded using the updateblastdb.pl script from BLAST+ package[76] and the kraken2 program [29], respectively. The fasta files of RefSeq fungal database was downloaded from the NCBI and converted to BLAST database using makblastdb command from the BLAST+ package[76], and was added to the kraken2 database library using kraken2 command [29]. We also build the standard kraken2 database for masking the contaminated regions within the fungal genomes using kraken2 command [29].

To generate the mock community database with only the species from the mock community, we downloaded the genomes of all species in the mock community from the NCBI according to their accessions (Supplementary Table S1), and concatenated them with the three newly assembled genomes of *Candida rugosa, Candida mesorugosa* and *Cryotococcus magnus.* Following the previous pipeline [77], we then performed a kraken2 search to identify the potential contaminated regions in the concatenated fasta, and masked those regions using bedtools [71]. We also masked the low complexity regions using the dustmasker from BLAST+ package [76]. To enable new genomes to be indexed by blastn, we updated the taxonomic map file by adding the fasta headers of the three new genomes and manually assigned their taxonomic ID in the file. Lastly, we used the makeblastdb program to construct the mock community database.

For amplicon data analysis, we used two versions of fungal ITS database from the NCBI and UNITE, plus the fungal 18S, 28S database from the NCBI. All of them are downloaded as fasta format in February 2020 and added to the kraken2 database library using kraken2 command [29].

### Data analysis

For Shotgun metagenomics datasets, we first used blastn (version 2.10.1) and kraken2 (version 2.0.8) to assign the NCBI taxonomic ID for each Illumina contig or Nanopore read. During the classification, we found one contamination species *Purpureocillium lilacinum* always present in all samples with a significant abundance (10-20%). Therefore, we added this species into the true species list. The best hit from BLAST or species with the highest k-mer counts for each read and/or contig was retained for further analysis. After classification, we used python pandas module to merge information from different output files, and used ete3 module [78] to assign taxonomic information to each read or contigs. The relative abundance of each classification were calculated based on the total length of Nanopore reads of total coverage of Illumina contigs. We used python numpy and math module for all statistical analysis.

For amplicon datasets, we sequenced each sample with three technical replicates. The classification workflow was different for datasets with different sequencing technologies. We only used QIME2 workflow plus the UNITE database for the Illumina amplicon data, since it is the only widely used method for classification. The paired end reads were denoised using the DADA2[79] plugin and assigned taxonomic information using the q2-feature-classifier [80] plugin. The QIME2 classifier was trained by the database sequence before classification. The classification output .qzv files were visualized by the QIME2 view website (https://view.qiime2.org/) and the feature-frequency csv file was extracted from the website. We then used python numpy and math module for the mathematical analysis and used seaborn module for generating figures.

For nanopore amplicon datasets, we used kraken2 as the k-mer based algorithm and minimap2 as the alignment based algorithm. The kraken2 command is the same as the kraken2 analysis for the shotgun metagenomics datasets, only using different databases. For the minimap2 analysis, we extracted the accessions of the best hits from the output files, and searched their corresponding taxonomic ID from the NCBI taxonomic map (downloaded from https://ftp.ncbi.nih.gov/pub/taxonomy/accession2taxid/nucl_wgs.accession2taxid.gz, in June 2020) using python pandas module. We then merge information from different output files, and used ete3 module again to assign taxonomic information to each read.

## Declarations

### Ethics approval and consent to participate

Not applicable.

### Consent for publication

Not applicable.

## Data Availability

All sequencing data was submitted to NCBI Short Read Archive (SRA) under the BioProject PRJNA725368 including eight accessions: SRX10705648, SRX10705649, SRX10705650, SRX10705651, SRX10705695, SRX10705696 and SRX10705697.

## Authors’ contributions

WM, ES, BS and JPR conceived the study and designed experiments. YH, LI and WTVH prepared the samples and generated sequencing data. YH, TE and AG performed the bioinformatics analysis. ES provided feedback on statistical analysis. All authors contributed to data analysis and manuscript writing. All authors read and approved the final manuscript.

## Acknowledgements

We thank Dr. Yu Lin for the insightful discussion and valuable suggestions to our manuscript. This work was supported by computational resources provided by the Australian Government through the National Computational Infrastructure (NCI) under the ANU Merit Allocation Scheme.

---

**Supplementary Table S1.** Metadata of the mock fungal community

**Supplementary Table S2.** Assembly statistics of the draft genomes of *Candida rugosa*, *Candida mesorugosa* and *Cryptococcus magnus* in the mock fungal community.

**Supplementary Figure S1.** Change of alignment metrics after applying cut-offs on Phred score.

