## Supplementary Table S2 for "Inferring species compositions of complex fungal communities from long- and short-read sequence data"

### Slide 1
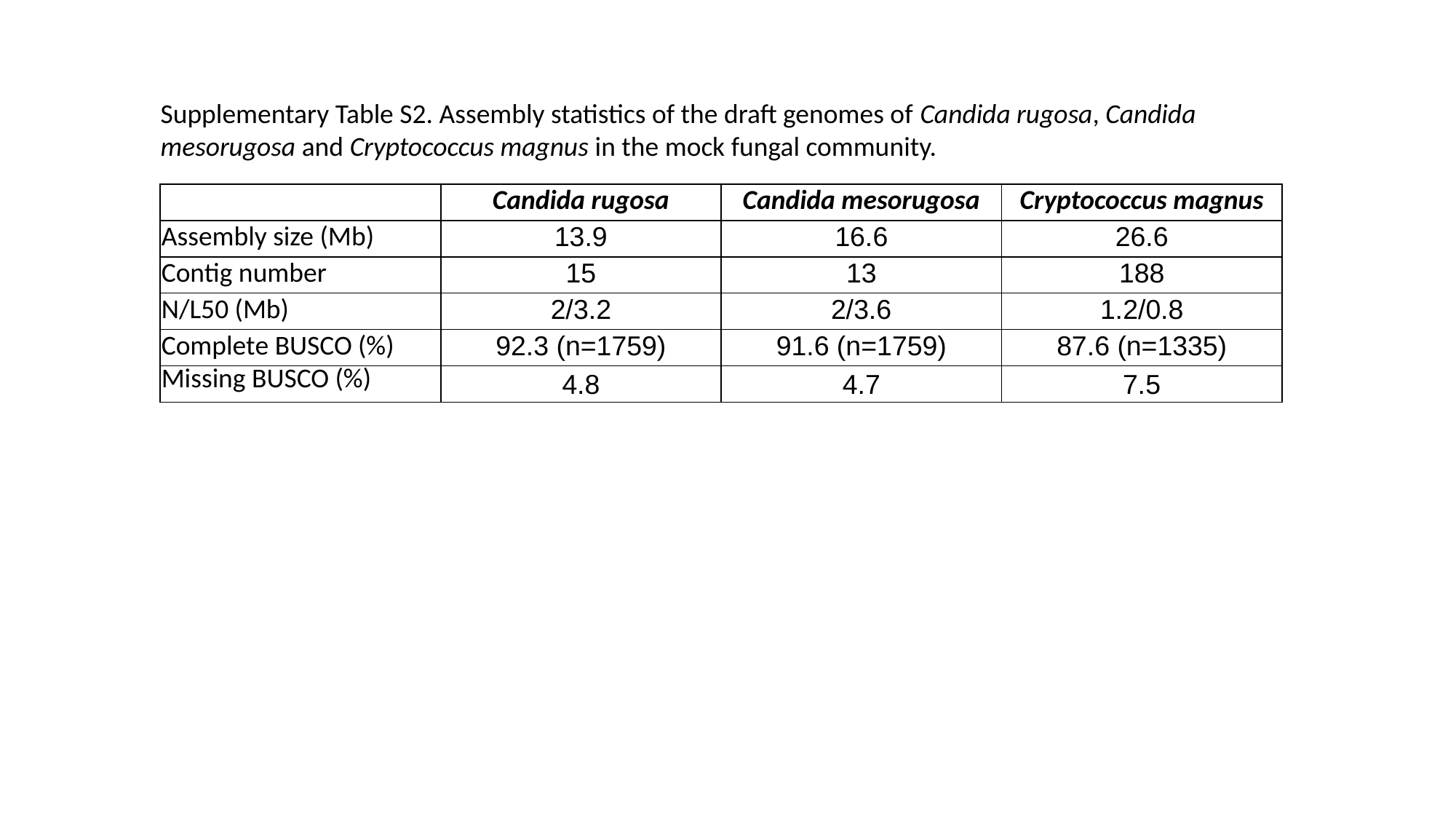

Supplementary Table S2. Assembly statistics of the draft genomes of Candida rugosa, Candida mesorugosa and Cryptococcus magnus in the mock fungal community.
| | Candida rugosa | Candida mesorugosa | Cryptococcus magnus |
| --- | --- | --- | --- |
| Assembly size (Mb) | 13.9 | 16.6 | 26.6 |
| Contig number | 15 | 13 | 188 |
| N/L50 (Mb) | 2/3.2 | 2/3.6 | 1.2/0.8 |
| Complete BUSCO (%) | 92.3 (n=1759) | 91.6 (n=1759) | 87.6 (n=1335) |
| Missing BUSCO (%) | 4.8 | 4.7 | 7.5 |
