## Supplementary Table S1 for "Inferring species compositions of complex fungal communities from long- and short-read sequence data"

### Slide 1
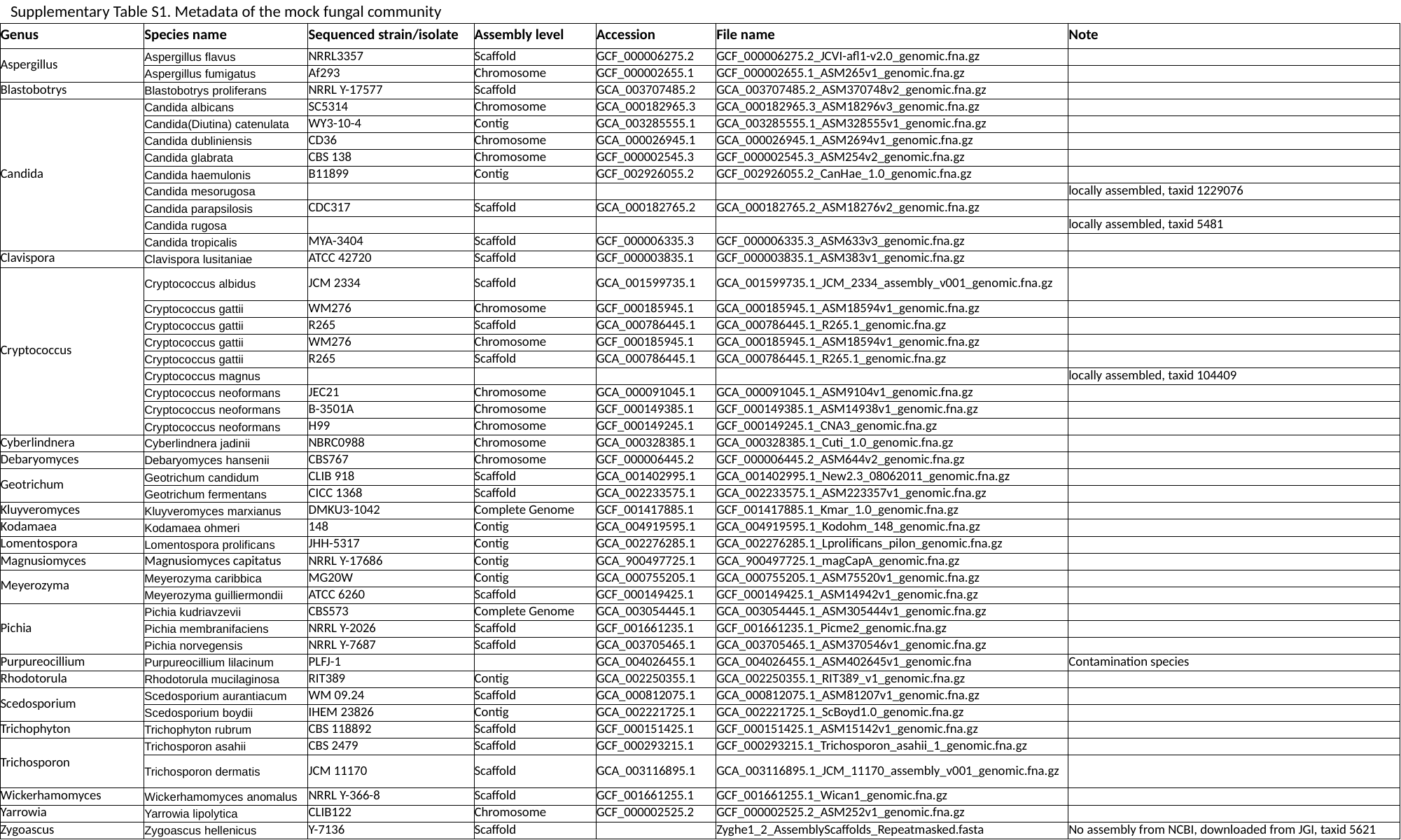

Supplementary Table S1. Metadata of the mock fungal community
| Genus | Species name | Sequenced strain/isolate | Assembly level | Accession | File name | Note |
| --- | --- | --- | --- | --- | --- | --- |
| Aspergillus | Aspergillus flavus | NRRL3357 | Scaffold | GCF\_000006275.2 | GCF\_000006275.2\_JCVI-afl1-v2.0\_genomic.fna.gz | |
| | Aspergillus fumigatus | Af293 | Chromosome | GCF\_000002655.1 | GCF\_000002655.1\_ASM265v1\_genomic.fna.gz | |
| Blastobotrys | Blastobotrys proliferans | NRRL Y-17577 | Scaffold | GCA\_003707485.2 | GCA\_003707485.2\_ASM370748v2\_genomic.fna.gz | |
| Candida | Candida albicans | SC5314 | Chromosome | GCA\_000182965.3 | GCA\_000182965.3\_ASM18296v3\_genomic.fna.gz | |
| | Candida(Diutina) catenulata | WY3-10-4 | Contig | GCA\_003285555.1 | GCA\_003285555.1\_ASM328555v1\_genomic.fna.gz | |
| | Candida dubliniensis | CD36 | Chromosome | GCA\_000026945.1 | GCA\_000026945.1\_ASM2694v1\_genomic.fna.gz | |
| | Candida glabrata | CBS 138 | Chromosome | GCF\_000002545.3 | GCF\_000002545.3\_ASM254v2\_genomic.fna.gz | |
| | Candida haemulonis | B11899 | Contig | GCF\_002926055.2 | GCF\_002926055.2\_CanHae\_1.0\_genomic.fna.gz | |
| | Candida mesorugosa | | | | | locally assembled, taxid 1229076 |
| | Candida parapsilosis | CDC317 | Scaffold | GCA\_000182765.2 | GCA\_000182765.2\_ASM18276v2\_genomic.fna.gz | |
| | Candida rugosa | | | | | locally assembled, taxid 5481 |
| | Candida tropicalis | MYA-3404 | Scaffold | GCF\_000006335.3 | GCF\_000006335.3\_ASM633v3\_genomic.fna.gz | |
| Clavispora | Clavispora lusitaniae | ATCC 42720 | Scaffold | GCF\_000003835.1 | GCF\_000003835.1\_ASM383v1\_genomic.fna.gz | |
| Cryptococcus | Cryptococcus albidus | JCM 2334 | Scaffold | GCA\_001599735.1 | GCA\_001599735.1\_JCM\_2334\_assembly\_v001\_genomic.fna.gz | |
| | Cryptococcus gattii | WM276 | Chromosome | GCF\_000185945.1 | GCA\_000185945.1\_ASM18594v1\_genomic.fna.gz | |
| | Cryptococcus gattii | R265 | Scaffold | GCA\_000786445.1 | GCA\_000786445.1\_R265.1\_genomic.fna.gz | |
| | Cryptococcus gattii | WM276 | Chromosome | GCF\_000185945.1 | GCA\_000185945.1\_ASM18594v1\_genomic.fna.gz | |
| | Cryptococcus gattii | R265 | Scaffold | GCA\_000786445.1 | GCA\_000786445.1\_R265.1\_genomic.fna.gz | |
| | Cryptococcus magnus | | | | | locally assembled, taxid 104409 |
| | Cryptococcus neoformans | JEC21 | Chromosome | GCA\_000091045.1 | GCA\_000091045.1\_ASM9104v1\_genomic.fna.gz | |
| | Cryptococcus neoformans | B-3501A | Chromosome | GCF\_000149385.1 | GCF\_000149385.1\_ASM14938v1\_genomic.fna.gz | |
| | Cryptococcus neoformans | H99 | Chromosome | GCF\_000149245.1 | GCF\_000149245.1\_CNA3\_genomic.fna.gz | |
| Cyberlindnera | Cyberlindnera jadinii | NBRC0988 | Chromosome | GCA\_000328385.1 | GCA\_000328385.1\_Cuti\_1.0\_genomic.fna.gz | |
| Debaryomyces | Debaryomyces hansenii | CBS767 | Chromosome | GCF\_000006445.2 | GCF\_000006445.2\_ASM644v2\_genomic.fna.gz | |
| Geotrichum | Geotrichum candidum | CLIB 918 | Scaffold | GCA\_001402995.1 | GCA\_001402995.1\_New2.3\_08062011\_genomic.fna.gz | |
| | Geotrichum fermentans | CICC 1368 | Scaffold | GCA\_002233575.1 | GCA\_002233575.1\_ASM223357v1\_genomic.fna.gz | |
| Kluyveromyces | Kluyveromyces marxianus | DMKU3-1042 | Complete Genome | GCF\_001417885.1 | GCF\_001417885.1\_Kmar\_1.0\_genomic.fna.gz | |
| Kodamaea | Kodamaea ohmeri | 148 | Contig | GCA\_004919595.1 | GCA\_004919595.1\_Kodohm\_148\_genomic.fna.gz | |
| Lomentospora | Lomentospora prolificans | JHH-5317 | Contig | GCA\_002276285.1 | GCA\_002276285.1\_Lprolificans\_pilon\_genomic.fna.gz | |
| Magnusiomyces | Magnusiomyces capitatus | NRRL Y-17686 | Contig | GCA\_900497725.1 | GCA\_900497725.1\_magCapA\_genomic.fna.gz | |
| Meyerozyma | Meyerozyma caribbica | MG20W | Contig | GCA\_000755205.1 | GCA\_000755205.1\_ASM75520v1\_genomic.fna.gz | |
| | Meyerozyma guilliermondii | ATCC 6260 | Scaffold | GCF\_000149425.1 | GCF\_000149425.1\_ASM14942v1\_genomic.fna.gz | |
| Pichia | Pichia kudriavzevii | CBS573 | Complete Genome | GCA\_003054445.1 | GCA\_003054445.1\_ASM305444v1\_genomic.fna.gz | |
| | Pichia membranifaciens | NRRL Y-2026 | Scaffold | GCF\_001661235.1 | GCF\_001661235.1\_Picme2\_genomic.fna.gz | |
| | Pichia norvegensis | NRRL Y-7687 | Scaffold | GCA\_003705465.1 | GCA\_003705465.1\_ASM370546v1\_genomic.fna.gz | |
| Purpureocillium | Purpureocillium lilacinum | PLFJ-1 | | GCA\_004026455.1 | GCA\_004026455.1\_ASM402645v1\_genomic.fna | Contamination species |
| Rhodotorula | Rhodotorula mucilaginosa | RIT389 | Contig | GCA\_002250355.1 | GCA\_002250355.1\_RIT389\_v1\_genomic.fna.gz | |
| Scedosporium | Scedosporium aurantiacum | WM 09.24 | Scaffold | GCA\_000812075.1 | GCA\_000812075.1\_ASM81207v1\_genomic.fna.gz | |
| | Scedosporium boydii | IHEM 23826 | Contig | GCA\_002221725.1 | GCA\_002221725.1\_ScBoyd1.0\_genomic.fna.gz | |
| Trichophyton | Trichophyton rubrum | CBS 118892 | Scaffold | GCF\_000151425.1 | GCF\_000151425.1\_ASM15142v1\_genomic.fna.gz | |
| Trichosporon | Trichosporon asahii | CBS 2479 | Scaffold | GCF\_000293215.1 | GCF\_000293215.1\_Trichosporon\_asahii\_1\_genomic.fna.gz | |
| | Trichosporon dermatis | JCM 11170 | Scaffold | GCA\_003116895.1 | GCA\_003116895.1\_JCM\_11170\_assembly\_v001\_genomic.fna.gz | |
| Wickerhamomyces | Wickerhamomyces anomalus | NRRL Y-366-8 | Scaffold | GCF\_001661255.1 | GCF\_001661255.1\_Wican1\_genomic.fna.gz | |
| Yarrowia | Yarrowia lipolytica | CLIB122 | Chromosome | GCF\_000002525.2 | GCF\_000002525.2\_ASM252v1\_genomic.fna.gz | |
| Zygoascus | Zygoascus hellenicus | Y-7136 | Scaffold | | Zyghe1\_2\_AssemblyScaffolds\_Repeatmasked.fasta | No assembly from NCBI, downloaded from JGI, taxid 5621 |
