## Supplementary figures and images for "Inferring species compositions of complex fungal communities from long- and short-read sequence data"

### FigureS1_qscore.png

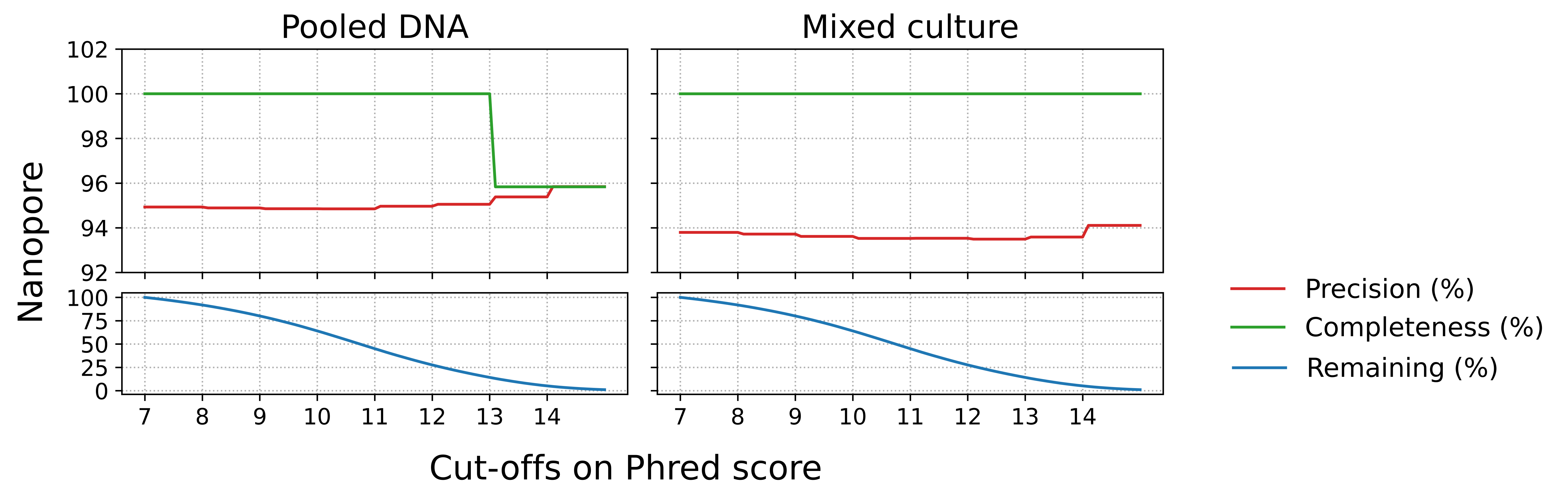
